# Chromatin architecture dictates epigenetic memory

**DOI:** 10.64898/2026.04.08.717143

**Authors:** Ziming Zhao, Jie Lin

**Affiliations:** Center for Quantitative Biology, Peking University, Beijing, China; Peking-Tsinghua Center for Life Sciences, Peking University, Beijing, China

## Abstract

Epigenetic memory encoded via histone modifications must remain robust across cell divisions. However, the physical rules governing how chromatin architecture regulates the maintenance and reorganization of epigenetic patterns remain unclear. Here, we develop a spreading-writing-erasing theory of epigenetic memory incorporating chromatin compartmentalization, enzyme limitation, and mark spreading. Our model demonstrates that self-sustaining epigenetic patterns emerge naturally without sequence insulators, while the establishment and erasure of heterochromatic compartments is governed by finite thresholds. Crucially, we uncover that the power-law exponent of chromatin contact probability dictates memory stability by buffering against spontaneous compartment collapse or expansion. Our theory also predicts progressive compartment fusion over cell generations, setting an upper limit of about 60 generations before senescence for human cells and explaining Hi-C map dynamics in senescent cells. Our theory also predicts a positive correlation between the contact-decay exponent and organismal complexity across species, supported by analysis of Hi-C maps across 873 species.

## INTRODUCTION

Epigenetic memory, often encoded as chemical modifications of DNA-bound histones, can be maintained across many cell generations and can also be erased during processes such as differentiation and induced reprogramming of differentiated cells [1–8]. While many details are known about how chemical modifications of histones correlate with gene expression [9–13], and how parental histones are segregated during DNA replication [14–16], fundamental questions remain unresolved. First, how do cells maintain epigenetic memory across cell cycles against the inevitable dilution of epigenetic marks by DNA replication? Second, how do cells modify epigenetic memory during reprogramming processes and differentiation? Third, how does large-scale chromatin architecture shape the stability and reorganization of epigenetic memory?

Chromosome conformation capture and imaging analyses have revealed that chromatin can be categorized into euchromatic and heterochromatic compartments with distinct epigenetic marks, known as A/B compartments. Genomic loci belonging to the same type of compartment are, on average, spatially closer than loci from different compartments [17–22]. Meanwhile, reader-writer enzymes spread epigenetic marks between spatially nearby histones [23–26]. The coupling between mark spreading and compartmentalization leads to a positive feed-back between them, which has been suggested to generate epigenetic memory, i.e., stable configurations of histone marks [27–32]. Recently, Owen et al. showed that stable epigenetic memory also requires a limited abundance of reader-writer enzymes [33].

However, the minimal conditions under which epigenetic patterns of histone modifications across the chromosome remain robustly stable across multiple cell cycles are not yet understood [33]. A mathematically tractable framework that jointly accounts for the maintenance, regulation, and long-term reorganization of epigenetic patterns is still lacking. Moreover, the implications of chromatin architecture for memory stability, its effect on senescence-associated chromatin reorganization, and the evolutionary changes in genome architecture across species remain elusive.

In this study, we develop a spreading-writing-erasing (SWE) theory of epigenetic memory, in which a one-dimensional, coarse-grained pattern of repressive histone marks represents epigenetic states. The model incorporates spreading, writing, and erasing of the epigenetic mark and generates self-sustaining epigenetic patterns across multiple cell generations. The theory further identifies finite thresholds of active writing and erasing for establishing and removing heterochromatic compartments, leading to history-dependent epigenetic patterns. Together, these results show that robust yet flexible epigenetic memory arises from the coupling between chromatin compartmentalization, contact-dependent mark spreading, and competition for limited reader-writer enzymes.

Our theory further predicts senescence-associated reorganization of epigenetic patterns, including compartment fusion and loss of fine-scale structures, consistent with Hi-C maps of human fibroblasts during replicative senescence. For human cells, the model predicts that epigenetic patterns can persist for up to approximately 60 generations before distinct reorganization, comparable to the replicative lifespan of primary human fibroblasts.

Our theory also unveils the critical role of chromatin spatial structure, quantified by the power-law exponent *n* of the contact probability between chromatin loci as a function of genomic distance. A larger *n*, corresponding to a faster decay of the contact probability, favors the stability of epigenetic memory and chromatin compartmentalization, reflected by the checkerboard pattern in the Hi-C maps. We verify our predictions using a comprehensive Hi-C dataset spanning 873 species [34] and indeed observe a positive correlation between the checkerboard score and the *n* exponent across species. More importantly, our theory suggests that *n* may positively correlate with organismal complexity, consistent with the data analysis and indicating a broader link between chromatin architecture and biological complexity.

Together, our work provides a physical framework for understanding how chromatin architecture enables epigenetic memory to keep robust while remaining flexible to regulation and gradually leading to cellular senescence. More broadly, the cross-species variation in chromatin contact scaling suggests an evolutionary link between genome architecture, epigenetic regulation, and organismal complexity.

## RESULTS

### A. Spreading-writing-erasing model generates robust epigenetic memory

We model the chromosome as a one-dimensional chain of total length *L*, with genomic coordinate *x* measured in 1-kb units. We encode epigenetic memory by an epigenetic mark level *m*(*x*) ∈ [0, 1], where euchromatic and heterochromatic loci correspond to low and high *m* values, respectively. One can consider *m*(*x*) as a coarsegrained variable that integrates the effects of multiple histone marks (e.g. H3K9me3, H3K27me3, etc), allowing us to study generic mechanisms underlying epigenetic memory. The dynamics of *m*(*x*) consist of spreading, erasing, and writing (Figure 1A):

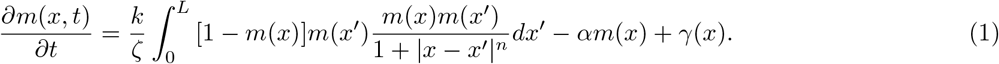

The first term on the right side of Equation 1 represents mark spreading catalyzed by the reader-writer enzyme, which copies the epigenetic marks to spatially nearby un-marked regions such that the spreading rate is proportional to the contact probability. We account for four important biological factors: (i) the contact probability averaged over all paired loci decays as a power law with an exponent *n* for large genomic distance, i.e., *P* (*s*) *~s*^*−n*^ where *s* = |*x −x*|^*′*^ is the genomic distance between two loci [17]; (ii) spreading of epigenetic marks between heterochromatic regions is more efficient due to compartmentalization, captured by the term *m*(*x*)*m*(*x*^*′*^); (iii) the epigenetic marks always spread from high-*m* regions to low-*m* regions, captured by the term [1 *−m*(*x*)]*m*(*x*^*′*^); (iv) the abundance of reader-writer enzymes is limited [33, 35], captured by the normalization factor *ζ*, the total number of marked-unmarked pairs where spreading may occur (Methods section 1). This normalization factor captures competition among spreading pairs for the limited reader-writer enzyme, which is critical for the stability of epigenetic memory as we show later. In writing Equation 1, we set the lower cutoff of the power-law decay in the contact probability as the length unit (1 kb) for simplicity, aligning with experiments [17].

The second term *αm*(*x*) represents the erasure of epigenetic marks, including passive erasing due to DNA replication and active erasing by an eraser enzyme (e.g., histone demethylases) [36–42]. In this work, we mostly model the dilution effect of DNA replication as continuous degradation unless otherwise mentioned. The last term, *γ*(*x*), represents active writing mediated by the writer enzyme (e.g., histone methyltransferases) [43]. For active erasing and writing, *α*(*x*) and *γ*(*x*) are sequence-dependent [11, 44–48]. We remark that the normalization factor *ζ* restricts the total mark level as∫ *m*(*x*)*dx* = *k/α* in the absence of active writing. We set the passive erasing rate *α*_0_ = 1, and the duration of one cell generation is therefore ln 2, unless otherwise mentioned. The derivation of the continuous SWE model from a discrete version is included in the Supplemental Material section 1.

We test whether the SWE model generates robust epigenetic memory and set the exponent *n* = 1.1 in the contact probability to simulate human cells [17]. The value of *n* is critical for epigenetic memory, as we show later. We initialize the system with several Gaussian packets of *m*(*x*), and it quickly self-organizes into stable compartments with high *m* values (Methods section 2). To focus on maintenance of epigenetic state (i.e., cell identity) against DNA replication here, active writing is not added. Intriguingly, the simulated epigenetic pattern persists over many generations (Figure 1B and Movie S1). The simulated Hi-C contact maps (Methods section 3) also reproduce the characteristic checkerboard pattern of A/B compartments (Figure 1C) [17, 18, 20, 22]. Notably, the compartments are self-organized with sharp boundaries and do not require insulator elements (e.g., CTCF), sequence-specific bookmarking, or fine-tuning of parameters, in contrast to several previous theoretical models [27, 29–31, 49]. Our model is consistent with experiments showing that A/B compartmentalization is preserved upon CTCF depletion [50].

**FIG. 1.**
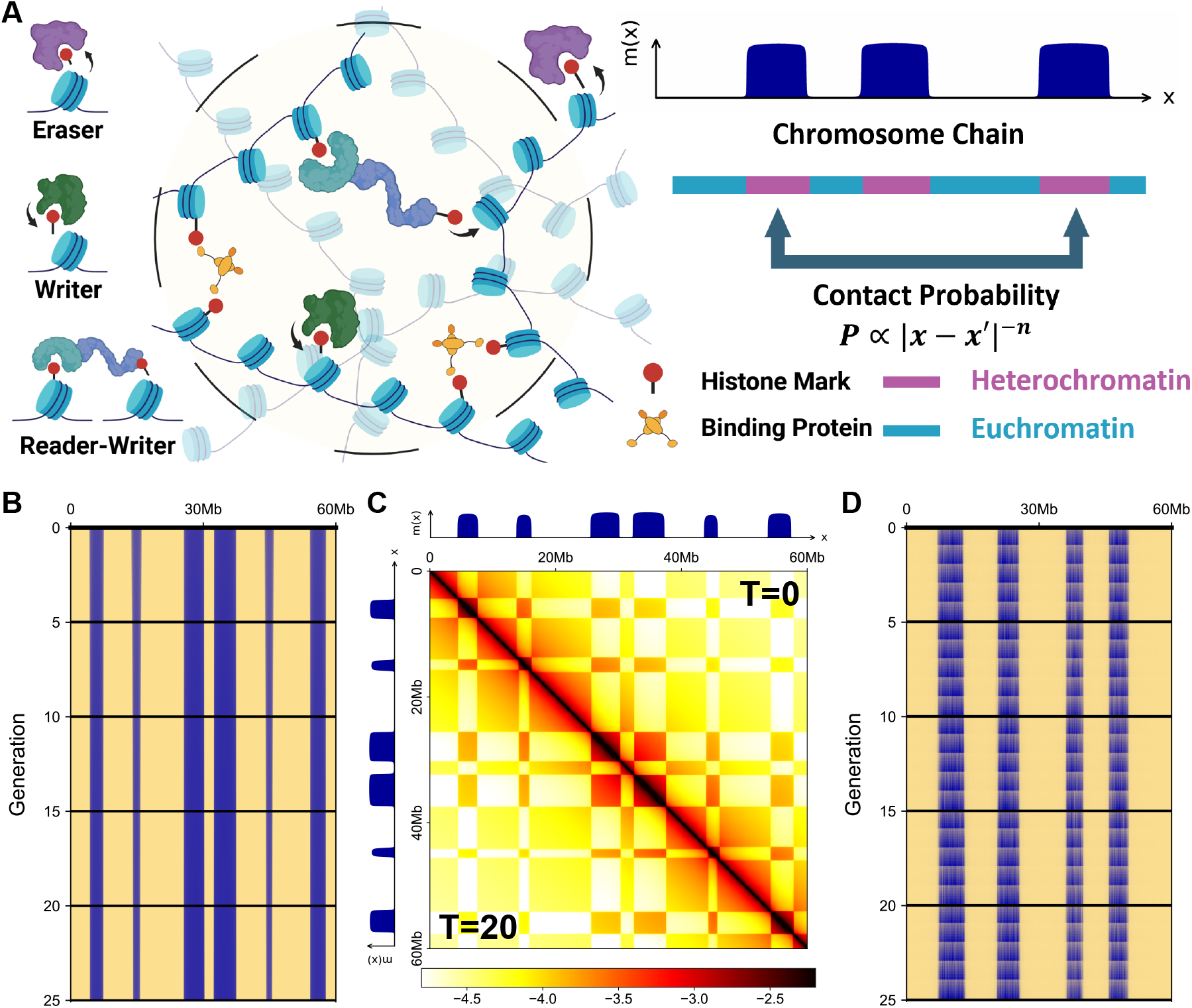
The spreading-writing-erasing model captures the stability of epigenetic memory. (A) Spreading of epigenetic marks is mediated by the reader-writer enzyme between spatially proximal histones. Writer and eraser enzymes add and remove marks at sequence-dependent loci. Binding proteins bridging nucleosomes with similar histone marks are also shown. Epigenetic memory is encoded in the histone mark level variable *m*(*x*) along the chromosome, with low and high values corresponding to euchromatin and heterochromatin. (B) Time evolution of the mark pattern through 25 generations with continuous degradation, where blue indicates a high mark level. (C) Comparison of simulated Hi-C contact maps at generation 0 (upper triangle) and 20 (lower triangle), showing persistent A/B compartmentalization, with the mark level *m*(*x*) shown alongside. (D) Time evolution of the mark pattern through 25 generations where stochastic segregation of histone marks due to periodic DNA replication is explicitly modeled.

We further test whether epigenetic memory is robust to stochasticity by modeling DNA replication as discrete events with random segregation of histone marks (Methods section 2). Again, the simulated epigenetic pattern is robust to abrupt and stochastic decreases in the mark level (Figure 1D), suggesting that the mathematical form of passive dilution in the model is not critical for memory maintenance. We highlight that the global competition for limited numbers of reader-writer enzymes is crucial for stabilizing epigenetic memory, as recently highlighted by Owen et al. [33]. To verify this idea, we simulate an alternative model with unlimited enzymes, and it fails to maintain stable compartments (Supplemental Material section 2). Compartments either vanish or expand boundlessly depending on the spreading rate (Figure S2). The unlimited-enzyme model also fails to maintain epigenetic memory against disturbance and periodic DNA replication (Figure S3).

### Bistability of epigenetic patterns and effects of chromatin architecture

We next seek to find the conditions for generating new compartments by nucleation, which can happen during cell-state transitions [11, 51–56]. We introduce a uniform active writing strength *γ* to a nucleation region with size *b*, inside which *γ*(*x*) *>* 0 and outside which *γ*(*x*) = 0.

By balancing the rates of spreading, writing, and erasing in Equation 1, the equilibrium mark level *m* that is approximately constant inside the compartments satisfies (Supplemental Material section 3),

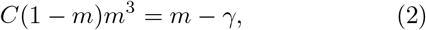

with the constant 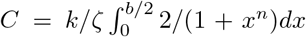. Equation 2 predicts that as the writing strength increases beyond threshold *γ*_*c*_, the induced mark level discontinuously jumps to a high-value state (Figure 2A), a consequence of the nonlinear and nonmonotonic spreading rate as a function of *m* (the left side of Equation 2 and Figure 2B). To verify the prediction, we simulate the SWE model and apply active writing to a nucleation region, successfully inducing a new compartment that remains self-sustaining after active writing is removed (Figure 2C). We remark that the induced compartment can expand beyond the nucleation region and also derive the growth criterion (the dashed line in Figure 2A; Supplemental Material section 3), which is always satisfied.

**FIG. 2.**
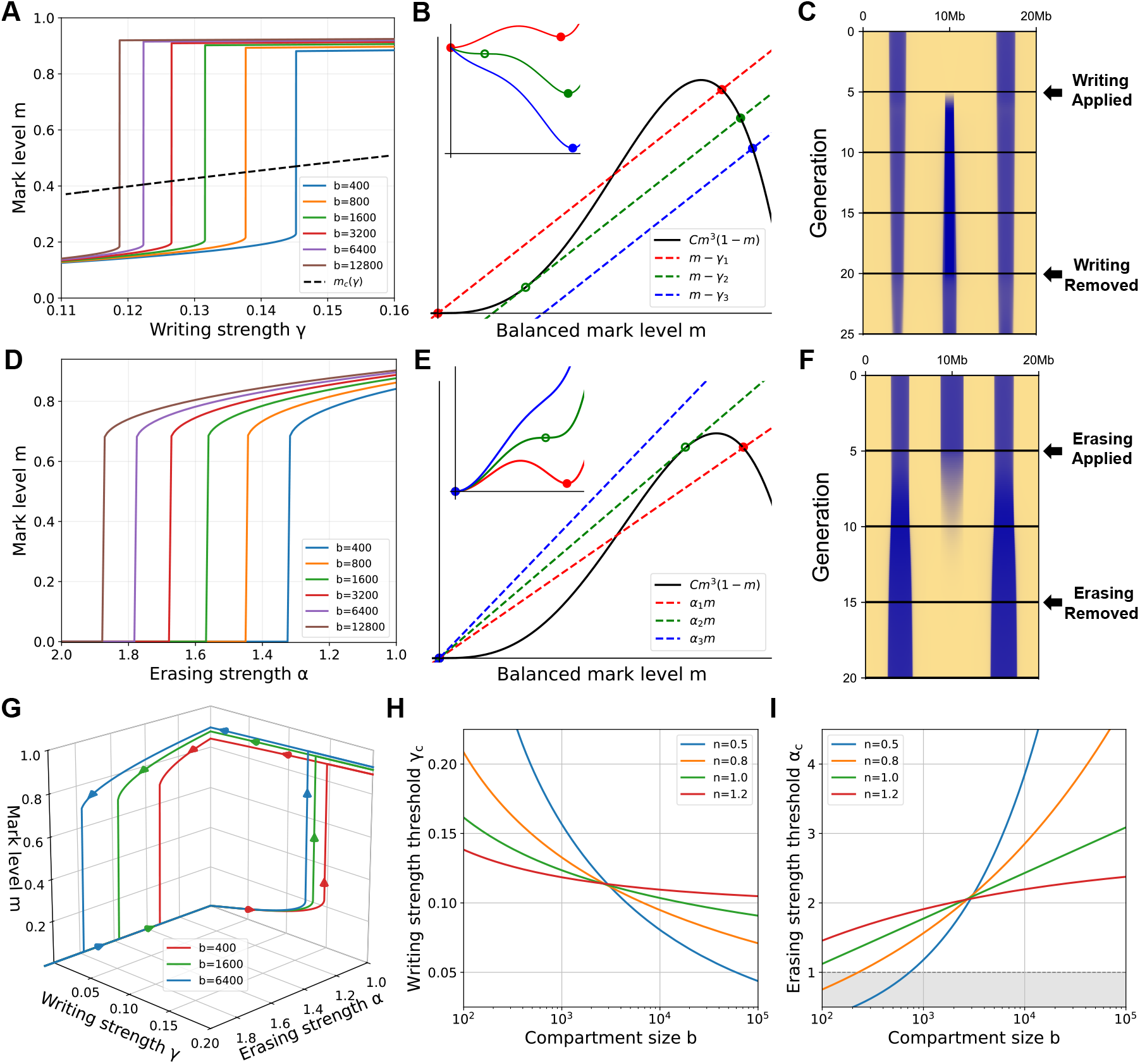
Origin of epigenetic memory and bistability. (A) The induced mark level *m* as a function of the active writing strength *γ*, with a given nucleation region size *b*. The dashed line is the threshold for the compartment to expand over the nucleation region. (B) Visualization of Equation 2, where the crossing point of the spreading rate and the net rate of writing and erasing determines the induced *m* level. Inset: the effective landscape showing the disappearance of the low *m* state as the writing strength exceeds the threshold *γ*_*c*_. (C) We apply local active writing to nucleate a new compartment at generation 5. The generated compartment grows and remains stable even after we remove active writing at generation 20. (D) The mark level *m* as a function of the erasing strength *α*. (E) Visualization of Equation 3 where the crossing point of the spreading and erasing rates determines the equilibrium *m* level. Inset: the effective landscape showing the disappearance of the high *m* state as the erasing strength exceeds the threshold *α*_*c*_. In (B) and (E), different dashed lines indicate different writing or erasing strengths, and the solid and open circles indicate stable and unstable fixed points, respectively. (F) We apply local active erasing to a compartment at generation 5. The compartment is erased and remains erased after active erasing is removed at generation 15. (G) The life cycle of a heterochromatic compartment as a function of the writing strength *γ* and erasing strength *α* with three nucleation region sizes *b*. (H) The minimum writing strength *γ*_*c*_ to establish a compartment as a function of the nucleation region size *b*. (I) The threshold erasing strength *α*_*c*_ to remove a compartment as a function of compartment size *b*. The shaded area highlights spontaneously unstable compartments with *α*_*c*_ below the passive dilution rate *α*_0_ = 1. Details of simulations related to this figure are in Methods section 2.

We next study the conditions for erasing a compartment by introducing a local erasing strength, *α >* 1, within an existing compartment. The balance between spreading and erasing determines the equilibrium mark level as (Supplemental Material section 3)

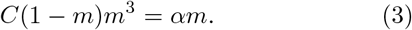

The balance condition predicts that as *α* increases, the mark level first decreases mildly and then rapidly drops to zero when the erasing strength exceeds a threshold *α*_*c*_ (Figure 2D and E). Equations 2 and 3 are numerically verified with model simulations (Figure S4, S5). Moreover, applying active erasing to an existing compartment is irreversible, as the removed compartment does not appear again after active erasing is removed (Figure 2F).

Taken together, the discontinuous dependence of the mark level on writing and erasing strengths gives rise to the hysteresis (i.e., history-dependent) life cycle of a heterochromatic compartment (Figure 2G), providing the physical basis of epigenetic memory. Our theory also suggests that cells do not need a high copy number of the writer or eraser enzyme during cell-state transition, since once compartments are formed or erased, the enzymes can be reallocated to other regions.

We study how the chromatin architecture, characterized by the exponent of the contact probability, affects epigenetic memory. We compute the threshold writing strength and erasing strength as a function of the nucleation size or compartment size *b* (Figure 2H, I). Interestingly, a large *n* exponent makes the thresholds less sensitive to the change in *b*, which we argue is critical for robust epigenetic memory: otherwise, large compartments can form spontaneously due to fluctuations in enzyme activity and small compartments can vanish spontaneously due to DNA replication (the shaded area in Figure 2I), which is detrimental to maintaining epigenetic memory. In the following, we show that chromatin architecture is indeed critical because it determines the duration of epigenetic-memory maintenance and how quickly a cell becomes senescent.

### Cellular senescence generated by chromatin reorganization

Hi-C maps have demonstrated loss of local chromatin connectivity and A/B compartment-switching during cellular senescence [57–61]. To determine whether our model captures chromatin reorganization during cellular senescence [61], we simulate the epigenetic pattern of a whole chromosome using the SWE model. Indeed, the simulated epigenetic patterns show a gradual loss of detailed structures (Figure 3A). In particular, nearby compartments can fuse, and large compartments gradually absorb smaller ones far away during chromatin reorganization, in agreement with the significant shift in domain boundaries and fusion of topologically associating domains (TADs) observed in senescent cells compared with proliferating cells [59].

**FIG. 3.**
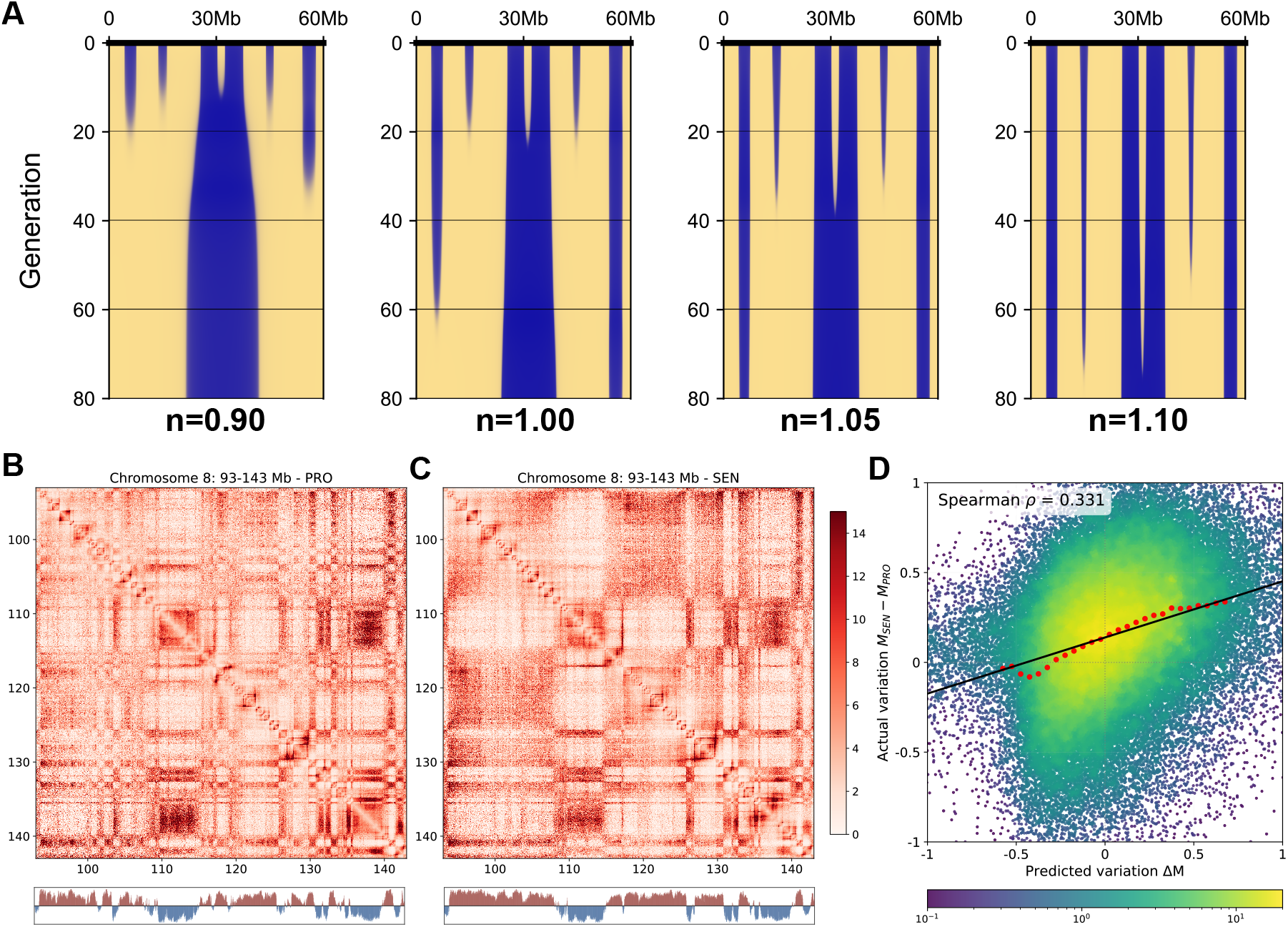
Cellular senescence due to chromatin reorganization. (A) Simulated evolution of the mark level with different exponents *n* given the same initial condition. Epigenetic memory lasts longer with a larger exponent *n*, and the detailed structures are finally lost. (B) Part of the Hi-C map of the human lung fibroblast cell in the proliferative state with the first principal component shown below. (C) The same as B, but for cells in the senescent state. (D) The model-predicted changes in the modification level Δ*m* vs. the changes inferred from the experimental data *m*_SEN_ *− m*_PRO_ between proliferative and senescent cells. Here, each data point corresponds to a genomic location. The red dots and the solid line represent the binned curve and linear regression of the distribution. The colorbar shows the relative data density.

Intriguingly, the larger the contact exponent *n*, the longer the epigenetic memory (i.e., cell identity) can be maintained (Figure 3A). In particular, for the simulation with *n* = 1.1, which is relevant for human cells, the maximum number of cell generations before significant reorganization of epigenetic patterns is around 60, approximately the number of cell divisions before primary human cells enter replicative senescence.

To quantify how maintenance time depends on compartment size and the contact exponent, we also study a simplified system in which the chromosome has two compartments of similar sizes. Interestingly, the maintenance time *t*_*m*_ increases as a power-law function of compartment size, and *t*_*m*_ becomes more sensitive to compartment size as *n* increases (Figure S6). These behaviors can be qualitatively understood from a simplified version of the SWE model where the dynamics are directly encoded by the compartment sizes (Supplemental Material section 5).

Next, we ask whether the SWE model can quantitatively explain the observed changes in Hi-C contact maps during cellular senescence by analyzing the Hi-C map of proliferative and senescent human fibroblast cells [62] (Figure 3B, C). Using the first principal component (PC1) of the Hi-C contact matrix as a proxy for the mark level *m*(*x*) (Methods section 4), we compute the modification change between the senescent and proliferative cells in two ways: the SWE model prediction Δ*m* derived from the proliferative state *m*_PRO_ alone, and the experimentally measured change *m*_SEN_*−m*_PRO_. Notably, the prediction correlates positively with the observed changes (Spearman’s *ρ* = 0.331, Figure 3D). Our analysis suggests that chromatin reorganization driven by mark spreading is an important mechanism of cellular senescence. Further, the model also predicts the changes between quiescent and senescent cells well (Spearman’s *ρ* = 0.376, Figure S7), while no correlation is observed for the changes between proliferative and quiescent cells (*ρ* = 0.033, Figure S8), suggesting that the quiescence-induced chromatin changes are not driven by the spontaneous compartment reorganization predicted by the SWE model.

### From chromatin architecture to organismal complexity

Next, we explore the implications of the SWE model from an evolutionary perspective. The SWE model predicts that if the contact probability decays slowly (i.e., with a small *n* exponent), the epigenetic pattern quickly loses its compartmental structure (Figure 3A). Regarding Hi-C contact maps, checkerboard patterns cannot be maintained. Therefore, we propose that the strength of the checkerboard patterns in Hi-C maps should correlate with the contact decay exponent *n* across different species.

To test this prediction, we analyze a comprehensive Hi-C dataset spanning 873 species across fungi and animals [34]. For each species, we compute the exponent *n* and the checkerboard score, a measure of compartmental organization (Methods section 4). Indeed, the checker-board score correlates positively with the contact exponent (Spearman’s *ρ* = 0.505, Figure 4A). Representative Hi-C maps illustrate that species with larger *n* exponents exhibit more pronounced checkerboard patterns in the Hi-C maps than species with smaller *n* (Figure 4B, C).

**FIG. 4.**
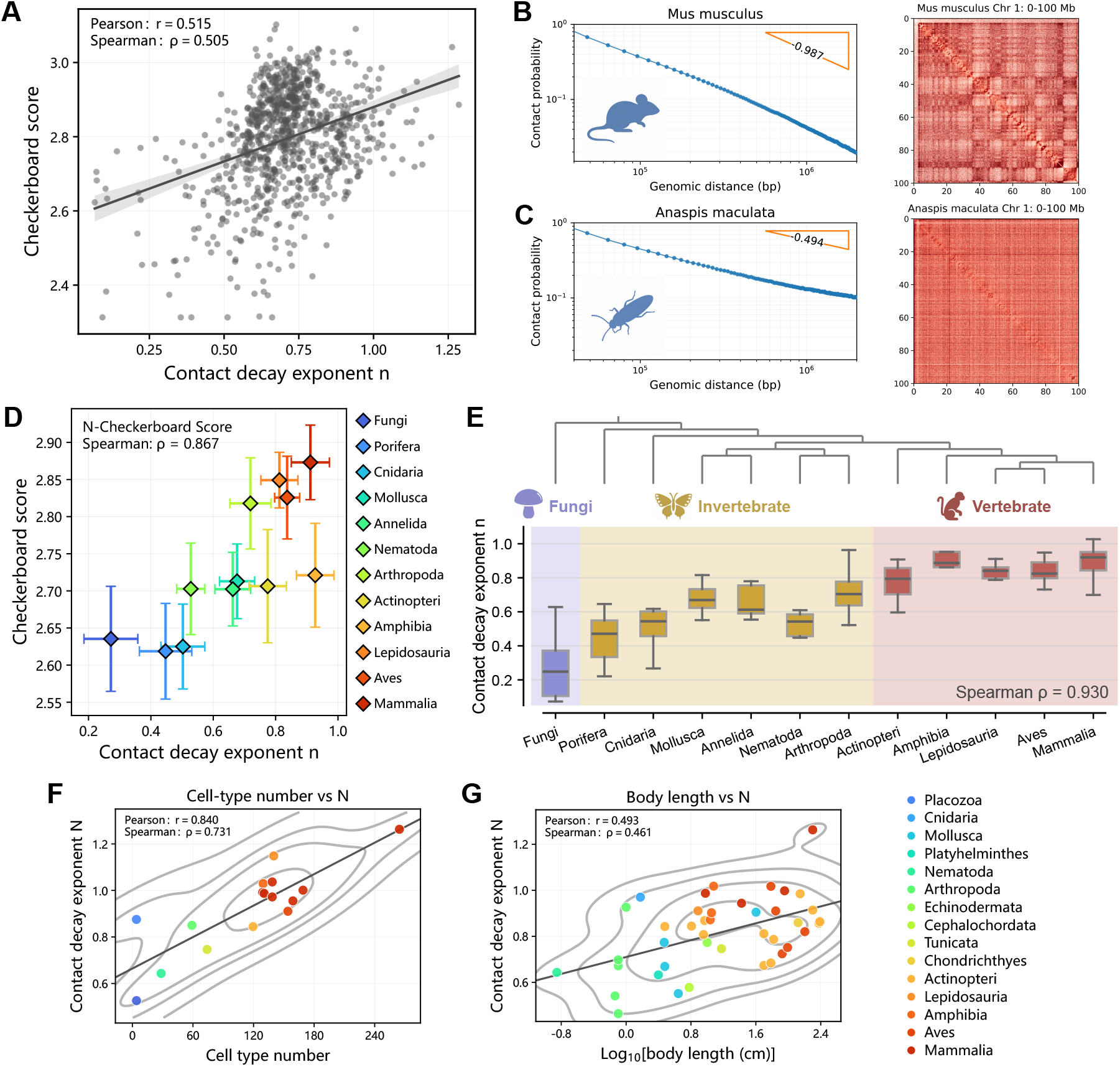
Evolutionary trend of chromatin architecture and organismal complexity. (A) Correlation between the contact decay exponent *n* and the checkerboard score across 873 species. Pearson and Spearman correlation coefficients are shown. (B, C) Contact probability as a function of genomic distance and the Hi-C maps of Chromosome 1 for *Mus musculus* (B, *n* = 0.987) and *Anaspis maculata* (C, *n* = 0.494), illustrating more pronounced compartmental organization in species with larger *n*. (D) Correlation between the contact decay exponent *n* and the checkerboard score across taxonomic groups. Different taxonomic groups are colored according to their phylogenetic order. (E) The evolutionary trend of the exponent *n* across phylogenetically ordered taxonomic groups. The Spearman correlation coefficient between the order of the taxonomic groups and the exponent *n* is shown. (F) Contact decay exponent *n* shows a positive relationship with the number of cell types. (G) Contact decay exponent *n* shows a positive relationship with body length. In (F) and (G), species are colored by phylogenetic order, and the solid lines represent linear regressions of the distributions.

We further hypothesize that more complex species require more diverse epigenetic regulation and therefore favor chromatin architectures that support more robust compartmentalization. Thus, we predict a positive correlation between organismal complexity and the exponent We confirm that as species are grouped by taxonomy, the contact exponent *n* correlates positively with the checkerboard score across taxonomic groups (Spearman’s *ρ* = 0.867, Figure 4D). More importantly, the contact exponent exhibits an increasing trend from fungi through invertebrates to vertebrates along the phylogenetic ordering (Spearman’s *ρ* = 0.930, Figure 4E), paralleling the evolutionary strengthening of checkerboard architecture [34].

We also test whether the exponent *n* correlates with organismal complexity directly, using a direct measure of complexity, the cell type number, as well as an indirect one, body length (Spearman’s *ρ* = 0.731 and 0.461 respectively, Figure 4F, G). Indeed, we find positive correlations between *n* and both parameters, suggesting that species with greater complexity tend to have a contact probability that decays faster.

## DISCUSSION

In this study, we introduce a minimal spreading-writing-erasing (SWE) model of epigenetic memory. The minimal ingredients of epigenetic memory include two critical mechanisms: attraction between heterochromatic loci and enzyme-mediated spreading of the epigenetic mark as the positive local feedback, and the limited abundance of the reader-writer enzyme as the negative global feedback, in agreement with Ref. [33]. These mechanisms stabilize epigenetic memory while permitting history-dependent switching between euchromatic and heterochromatic states under sufficiently strong regulation (Figure 2G).

An important prediction of the SWE model is that the scaling of chromatin contact probability modulates the stability of epigenetic memory. A large exponent *n* makes the establishment and removal of compartments less sensitive to domain size (Figure 2H, I) and leads to a longer duration of epigenetic memory (Figure 3A). The SWE model further predicts progressive reorganization of epigenetic patterns, characterized by compartment fusion and loss of fine-scale structures. Consistent with simulations, large-scale chromatin reorganizations are observed in the transition between proliferative and senescent cells (Figure 3B, C). The predicted changes in contact maps from the SWE model positively correlate with the experimental measurements, suggesting that chromatin reorganization is an important source of senescence.

The SWE model also predicts a positive correlation between the checkerboard score of Hi-C maps and the exponent *n*, consistent with our analysis of the Hi-C dataset across 873 species (Figure 4A-C). Furthermore, we propose that the *n* exponent should correlate with organismal complexity, as more complex species require more complex epigenetic regulation, reflected in the checker-board pattern in the Hi-C maps. Indeed, the *n* exponent increases as organismal complexity increases across fungi and animals. We also confirm this idea by directly computing the *n* exponent and measures of organismal complexity (e.g., cell type number). These observations suggest that chromatin architecture are probably under evolutionary selection to generate multiple cell types in multicellular organisms.

We highlight the coarse-grained nature of the SWE model, which neglects factors such as DNA methylation and fine-scale structures like chromatin loops. These simplifications allow analysis of the generic principles of epigenetic memory but limit system-specific predictions. As an application of our theory, we apply the SWE model to study how noise affects reprogramming efficiency of induced pluripotent stem cells. We find that increasing noise in parental histone segregation and shortening the doubling time can both enhance reprogramming efficiency (Supplemental Material section 6 and Figure S9), providing experimentally testable and potentially useful predictions. Future work could extend the model by introducing these specific regulatory factors and detailed modification kinetics. It would also be promising to apply the SWE model to dynamical data of Hi-C maps and epigenetic modifications across multiple cell generations.

## METHODS

### 1. The normalization factor in the SWE model

To take account of the limited abundance of the reader-writer enzyme, we introduce a normalization factor *ζ* in Equation 1,

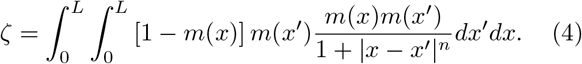

In the absence of active writing, integrating Equation 1 where *M* = ∫*m*(*x*)*dx*. Therefore, the total mark level is over *x* under a stable state leads to *dM/dt* = *k−αM* = 0, virtually constant and satisfies ∫*m*(*x*)*dx* = *k/α*.

### 2. Details of numerical simulations

For all simulations in this work, we initially introduce several Gaussian packets of *m*(*x*). We start recording data after 5 generations (*t* = 5 ln 2), as the packets rapidly self-organize into stable compartments with high *m* values, corresponding to generation 0. Details of each figure are listed as follows.

In Figure 1B, we set 6 initial Gaussian packets with different widths and locations and obtain 6 stable compartments from generation 0 to 25. In all the other similar figures, the initial number of compartments is also set by the initial number of Gaussian packets. In Figure 1D, we set the mark level at each site *x* to zero with probability *p* = 0.5 at every generation. In the meantime, we also include continuous dilution with a basal dilution rate *α*_*b*_ = 0.3.

In Figure 2A and D, we set *k/ζ* = 1.1. In Figure 2C, we apply *γ* = 0.9 to a nucleation region with size *b* = 1 Mb at generation 5. The compartment widens to *b≈* 1.5 Mb before active writing is removed at generation In Figure 2F, we increase the erasing rate to *α* = 1.5 in the central compartment and let it return to the passive dilution value at generation 15. In both panels, we set *k* = 5000 and *L* = 20000. In Figure 2H and I, the factor *k/ζ* is set by considering a chromosome composed of multiple compartments with a typical size *b*_0_ = 10^3.5^ (Supplemental Material section 4).

### 3. Hi-C map construction

In Figure 1, the simulated Hi-C map is generated by assuming the contact probability between two loci at *x*_1_ and *x*_2_ has the following form:

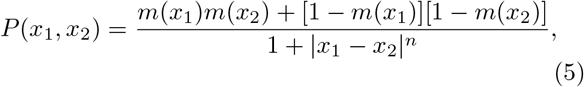

which captures the attraction between compartments of the same type, i.e., euchromatin (low *m*) and heterochromatin (high *m*).

### 4. Experimental data analysis

In Figure 3, Hi-C data for proliferating (PRO), quiescent (QUI), and replicatively senescent (SEN) LF1 human lung fibroblasts are from Ref. [62]. In Figure 3B and C, we extract PC1 from the Hi-C contact matrix of the PRO and SEN states, and compare the data with the compartment annotations to ensure that euchromatin corresponds to positive PC1 values.

In Figure 3D, to analyze the data in the SWE framework, we normalize the PC1 vector into 0 *< m*_*i*_ *<* 1, and compare the experimentally inferred modification change *m*_SEN_ *−m*_PRO_ with the predicted variation Δ*m* calculated as:

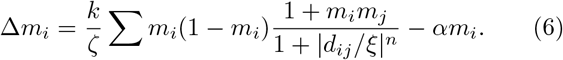

As in Equation 1, we set *k/ζ* = 0.9, *n* = 1.1, *α* = 1 and *ξ* = 1 kb in the prediction. We apply the same calculations to quiescence-to-senescent and proliferating-to-quiescence transitions (Figures S7 and S8).

In Figure 4, we use the cross-species Hi-C dataset from Ref. [34], which contains normalized Hi-C maps and checkerboard scores for species across fungi and animals. We exclude plants, which generally exhibit weak compartmentalization, as well as species with chromosomes too small to support robust analysis. After these exclusions, 873 species are retained for the cross-species analysis. For each species, we calculate the contact probability *P* (*s*) as a function of genomic distance *s* by averaging Hi-C contacts between loci separated by *s*. The contact-decay exponent *n* is estimated from the long-range regime according to

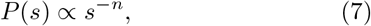

by linear regression of ln *P* vs. ln *s*.

In Figure 4A, to better assess the inter-taxon association between the contact decay exponent and checker-board score, we subsample the overrepresented arthropod species (*n*_arth_ = 664 in *n*_0_ = 873) to 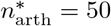. We obtain Pearson and Spearman correlation coefficients in 1000 resample replications as *r* = 0.5154*±* 0.0103 and *ρ* = 0.5045*±* 0.0148 (mean *±*standard deviation), respectively.

In Figure 4D and E, we group species into major taxonomic groups following the classification used by Ref. [34]. Taxonomic relationships are based on the NCBI taxonomy and arranged according to their phylogenetic positions. For each taxonomic group, the contact probability and checkerboard score are represented by the mean value. In Figure 4F and G, the cell type number and body length are from Ref. [34] and Ref. [63]. Body lengths are log_10_transformed before analysis.

## Supplemental material

Supplemental material is available for this paper.

## Funding

This work is supported by the National Natural Science Foundation of China (Grant No. 12474190), the National Key Research and Development Program of China (2024YFA0919600), and grants from the Peking-Tsinghua Center for Life Sciences.

## Competing interests

The authors declare no competing interests.

## Availability

The datasets and computer codes for the numerical simulations generated and analyzed during the current study are available from the corresponding author on reasonable request.

## Author contributions

Z.Z. and J.L. conceive the study and formulated the theoretical model. Z.Z. performs the numerical simulations and analyzed the data. All authors contributed to writing and reviewing the manuscript.

## Notes

### Competing Interest Statement

The authors have declared no competing interest.

### Summary of Updates

We added new analysis of Hi-C contact maps to the updated version.

## References

[1] K. Takahashi and S. Yamanaka, Induction of pluripotent stem cells from mouse embryonic and adult fibroblast cultures by defined factors, Cell 126, 663 (2006).

[2] I. B. Dodd, M. A. Micheelsen, K. Sneppen, and G. Thon, Theoretical analysis of epigenetic cell memory by nucleosome modification, Cell 129, 813 (2007).

[3] A. V. Probst, E. Dunleavy, and G. Almouzni, Epigenetic inheritance during the cell cycle, Nature reviews Molecular cell biology 10, 192 (2009).

[4] Y. Hirabayashi and Y. Gotoh, Epigenetic control of neural precursor cell fate during development, Nature reviews. Neuroscience 11, 377 (2010).

[5] R. D. Hawkins, G. C. Hon, L. K. Lee, Q. Ngo, R. Lister, M. Pelizzola, L. E. Edsall, S. Kuan, Y. Luu, S. Klugman, J. Antosiewicz-Bourget, Z. Ye, C. Espinoza, S. Agarwahl, L. Shen, V. Ruotti, W. Wang, R. Stewart, J. A. Thomson, J. R. Ecker, and B. Ren, Distinct epigenomic landscapes of pluripotent and lineage-committed human cells, Cell Stem Cell 6, 479 (2010).

[6] A. Angel, J. Song, C. Dean, and M. Howard, A polycomb-based switch underlying quantitative epigenetic memory, Nature 476, 105 (2011).

[7] K. Ragunathan, G. Jih, and D. Moazed, Epigenetic inheritance uncoupled from sequence-specific recruitment, Science 348, 1258699 (2015).

[8] Y. Atlasi and H. Stunnenberg, The interplay of epigenetic marks during stem cell differentiation and development, Nature Reviews Genetics 18 (2017).

[9] S. I. Grewal and D. Moazed, Heterochromatin and epigenetic control of gene expression, science 301, 798 (2003).

[10] R. Bonasio, S. Tu, and D. Reinberg, Molecular signals of epigenetic states, science 330, 612 (2010).

[11] N. Hathaway, O. Bell, C. Hodges, E. Miller, D. Neel, and G. Crabtree, Dynamics and memory of heterochromatin in living cells, Cell 149, 1447 (2012).

[12] S. Berry, M. Hartley, T. S. G. Olsson, C. Dean, and M. Howard, Local chromatin environment of a polycomb target gene instructs its own epigenetic inheritance, eLife 4, e07205 (2015).

[13] R. C. Allshire and H. D. Madhani, Ten principles of heterochromatin formation and function, Nat Rev Mol Cell Biol 19, 229 (2018).

[14] S. Liu, Z. Xu, H. Leng, P. Zheng, J. Yang, K. Chen, J. Feng, and Q. Li, Rpa binds histone h3-h4 and functions in dna replication–coupled nucleosome assembly, Science 355, 415 (2017).

[15] N. Reverón-Gómez, C. González-Aguilera, K. R. Stewart-Morgan, N. Petryk, V. Flury, S. Graziano, J. V. Johansen, J. S. Jakobsen, C. Alabert, and A. Groth, Accurate recycling of parental histones reproduces the histone modification landscape during dna replication, Molecular Cell 72, 239 (2018).

[16] T. M. Escobar, O. Oksuz, R. Saldana-Meyer, N. Descostes, R. Bonasio, and D. Reinberg, Active and repressed chromatin domains exhibit distinct nucleosome segregation during dna replication, Cell 179, 953 (2019).

[17] E. Lieberman-Aiden, N. L. van Berkum, L. Williams, M. Imakaev, T. Ragoczy, A. Telling, I. Amit, B. R. Lajoie, P. J. Sabo, M. O. Dorschner, R. Sandstrom, B. Bernstein, M. A. Bender, M. Groudine, A. Gnirke, J. Stamatoyannopoulos, L. A. Mirny, E. S. Lander, and J. Dekker, Comprehensive mapping of long-range interactions reveals folding principles of the human genome, Science 326, 289 (2009).

[18] S. Rao, M. Huntley, N. Durand, E. Stamenova, Bochkov, J. Robinson, A. Sanborn, I. Machol, A. Omer, E. Lander, and E. Aiden, A 3d map of the human genome at kilobase resolution reveals principles of chromatin looping, Cell 159, 1665 (2014).

[19] S. Wang, J.-H. Su, B. J. Beliveau, B. Bintu, J. R. Moffitt, C. ting Wu, and X. Zhuang, Spatial organization of chromatin domains and compartments in single chromosomes, Science 353, 598 (2016).

[20] M. J. Rowley and V. G. Corces, Organizational principles of 3d genome architecture, Nature Reviews Genetics 19, 789 (2018).

[21] M. Falk, Y. Feodorova, N. Naumova, M. Imakaev, B. R. Lajoie, H. Leonhardt, B. Joffe, J. Dekker, G. Fudenberg, and I. Solovei, Heterochromatin drives compartmentalization of inverted and conventional nuclei, Nature 570, 395 (2019).

[22] E. M. Hildebrand and J. Dekker, Mechanisms and functions of chromosome compartmentalization, Trends in Biochemical Sciences 45, 385 (2020).

[23] R. Margueron, N. Justin, K. Ohno, M. L. Sharpe, J. Son, r Drury, W. J., P. Voigt, S. R. Martin, W. R. Taylor, V. De Marco, V. Pirrotta, D. Reinberg, and S. J. Gamblin, Role of the polycomb protein eed in the propagation of repressive histone marks, Nature 461, 762 (2009).

[24] M. M. Muller, B. Fierz, L. Bittova, G. Liszczak, and T. W. Muir, A two-state activation mechanism controls the histone methyltransferase suv39h1, Nat Chem Biol 12, 188 (2016).

[25] O. Oksuz, V. Narendra, C.-H. Lee, N. Descostes, G. LeRoy, R. Raviram, L. Blumenberg, K. Karch, P. P. Rocha, B. A. Garcia, J. A. Skok, and D. Reinberg, Capturing the onset of prc2-mediated repressive domain formation, Molecular Cell 70, 1149 (2018).

[26] K. Kraft, K. E. Yost, S. E. Murphy, A. Magg, Y. Long, M. R. Corces, J. M. Granja, L. Wittler, S. Mundlos, T. R. Cech, A. N. Boettiger, and H. Y. Chang, Polycomb-mediated genome architecture enables long-range spreading of h3k27 methylation, Proceedings of the National Academy of Sciences 119, e2201883119 (2022).

[27] F. Erdel and E. C. Greene, Generalized nucleation and looping model for epigenetic memory of histone modifications, Proc Natl Acad Sci USA 113, E4180 (2016).

[28] D. Michieletto, E. Orlandini, and D. Marenduzzo, Polymer model with epigenetic recoloring reveals a pathway for the de novo establishment and 3d organization of chromatin domains, Physical Review X 6, 041047 (2016).

[29] D. Jost and C. Vaillant, Epigenomics in 3d: importance of long-range spreading and specific interactions in epigenomic maintenance, Nucleic Acids Res 46, 2252 (2018).

[30] S. H. Sandholtz, Q. MacPherson, and A. J. Spakowitz, Physical modeling of the heritability and maintenance of epigenetic modifications, Proc Natl Acad Sci U S A 117, 20423 (2020).

[31] M. Katava, G. Shi, and D. Thirumalai, Chromatin dynamics controls epigenetic domain formation, Biophys J 121, 2895 (2022).

[32] H. Zhang, X.-J. Tian, A. Mukhopadhyay, K. Kim, and J. Xing, Statistical mechanics model for the dynamics of collective epigenetic histone modification, Physical review letters 112, 068101 (2014).

[33] J. A. Owen, D. Osmanovic, and L. Mirny, Design principles of 3d epigenetic memory systems, Science 382, eadg3053 (2023).

[34] Y. Che, S. J. Bush, H. Lin, M. Li, X. Yang, Q. Xie, Y. Liu, D. Meng, and K. Ye, The evolution of high-order genome architecture revealed from 1,000 species, Cell 189, 3760 (2026).

[35] V. Ramesh, F. Liu, M. S. Minto, U. Chan, and A. E. West, Bidirectional regulation of postmitotic h3k27me3 distributions underlie cerebellar granule neuron maturation dynamics, Elife 12, e86273 (2023).

[36] Y. Shi, F. Lan, C. Matson, P. Mulligan, J. R. Whetstine, P. A. Cole, R. A. Casero, and Y. Shi, Histone demethylation mediated by the nuclear amine oxidase homolog lsd1, Cell 119, 941 (2004).

[37] M. Lee, C. Wynder, N. Cooch, and R. Shiekhattar, An essential role for corest in nucleosomal histone 3 lysine 4 demethylation, Nature 437, 432 (2005).

[38] Y.-i. Tsukada, J. Fang, H. Erdjument-Bromage, M. E. Warren, C. H. Borchers, P. Tempst, and Y. Zhang, Histone demethylation by a family of jmjc domaincontaining proteins, Nature 439, 811 (2006).

[39] N. P. Blackledge, J. C. Zhou, M. Y. Tolstorukov, A. M. Farcas, P. J. Park, and R. J. Klose, Cpg islands recruit a histone h3 lysine 36 demethylase, Molecular Cell 38, 179 (2010).

[40] N. Mosammaparast and Y. Shi, Reversal of histone methylation: Biochemical and molecular mechanisms of histone demethylases, Annu Rev Biochem 79, 155 (2010).

[41] S. M. Kooistra and K. Helin, Molecular mechanisms and potential functions of histone demethylases, Nat Rev Mol Cell Biol 13, 297 (2012).

[42] L. Peng, X. Li, H. Yang, H. Chen, Y. Yang, and S. Peng, Discovery and structural studies of histone demethylases, Frontiers in Epigenetics and Epigenomics 3, 1594400 (2025).

[43] S. Rea, F. Eisenhaber, D. O’Carroll, B. D. Strahl, Z.-W. Sun, M. Schmid, S. Opravil, K. Mechtler, C. P. Ponting, C. D. Allis, et al., Regulation of chromatin structure by site-specific histone h3 methyltransferases, Nature 406, 593 (2000).

[44] M.-C. Tsai, O. Manor, Y. Wan, N. Mosammaparast, J. K. Wang, F. Lan, Y. Shi, E. Segal, and H. Y. Chang, Long noncoding rna as modular scaffold of histone modification complexes, Science 329, 689 (2010).

[45] C. Beisel and R. Paro, Silencing chromatin: comparing modes and mechanisms, Nature Reviews Genetics 12, 123 (2011).

[46] A. J. Bannister and T. Kouzarides, Regulation of chromatin by histone modifications, Cell research 21, 381 (2011).

[47] M. Park, N. Patel, A. J. Keung, and A. S. Khalil, Engineering epigenetic regulation using synthetic read-write modules, Cell 176, 227 (2019).

[48] A. Tatarakis, H. Saini, J. Yu, W. Feng, C. A. Pinzon-Arteaga, and D. Moazed, Requirements for establishment and epigenetic stability of mammalian heterochromatin, Molecular Cell 85, 3388 (2025).

[49] D. Michieletto, M. Chiang, D. Colì, A. Papantonis, E. Orlandini, P. R. Cook, and D. Marenduzzo, Shaping epigenetic memory via genomic bookmarking, Nucleic Acids Research 46, 83 (2017).

[50] E. P. Nora, A. Goloborodko, A.-L. Valton, J. H. Gibcus, Uebersohn, N. Abdennur, J. Dekker, L. A. Mirny, and G. Bruneau, Targeted degradation of ctcf decouples local insulation of chromosome domains from genomic compartmentalization, Cell 169, 930 (2017).

[51] T. Yamada, W. Fischle, T. Sugiyama, C. D. Allis, and S. I. Grewal, The nucleation and maintenance of heterochromatin by a histone deacetylase in fission yeast, Molecular Cell 20, 173 (2005).

[52] C. Hodges and G. R. Crabtree, Dynamics of inherently bounded histone modification domains, Proc Natl Acad Sci U S A 109, 13296 (2012).

[53] L. Bintu, J. Yong, Y. E. Antebi, K. McCue, Y. Kazuki, N. Uno, M. Oshimura, and M. B. Elowitz, Dynamics of epigenetic regulation at the single-cell level, Science 351, 720 (2016).

[54] M. Obersriebnig, E. Pallesen, K. Sneppen, A. Trusina, and G. Thon, Nucleation and spreading of a heterochromatic domain in fission yeast, Nature communications 7, 11518 (2016).

[55] F. Laprell, K. Finkl, and J. Müller, Propagation of polycomb-repressed chromatin requires sequence-specific recruitment to dna, Science 356, 85 (2017).

[56] S. I. Grewal, The molecular basis of heterochromatin assembly and epigenetic inheritance, Molecular Cell 83, 1767 (2023).

[57] T. Chandra, P. Ewels, S. Schoenfelder, M. Furlan-Magaril, S. Wingett, K. Kirschner, J.-Y. Thuret, S. Andrews, P. Fraser, and W. Reik, Global reorganization of the nuclear landscape in senescent cells, Cell Reports 10, 471 (2015).

[58] S. W. Criscione, M. D. Cecco, B. Siranosian, Y. Zhang, J. A. Kreiling, J. M. Sedivy, and N. Neretti, Reorganization of chromosome architecture in replicative cellular senescence, Science Advances 2, e1500882 (2016).

[59] A. Zirkel, M. Nikolic, K. Sofiadis, J.-P. Mallm, C. A. Brackley, H. Gothe, O. Drechsel, C. Becker, J. Altmüller, N. Josipovic, T. Georgomanolis, L. Brant, J. Franzen, M. Koker, E. G. Gusmao, I. G. Costa, R. T. Ullrich, W. Wagner, V. Roukos, P. Nürnberg, D. Marenduzzo, K. Rippe, and A. Papantonis, Hmgb2 loss upon senescence entry disrupts genomic organization and induces ctcf clustering across cell types, Molecular Cell 70, 730 (2018).

[60] X. Zhang, X. Liu, Z. Du, L. Wei, H. Fang, Q. Dong, J. Niu, Y. Li, J. Gao, M. Zhang, W. Xie, and X. Wang, The loss of heterochromatin is associated with multiscale three-dimensional genome reorganization and aberrant transcription during cellular senescence, Genome Research 31 (2021).

[61] I. Olan and M. Narita, Senescence: An identity crisis originating from deep within the nucleus, Annual Review of Cell and Developmental Biology 38, 219 (2022).

[62] A. Dalgarno, S. A. Evans, M. M. Kelsey, T. A. Nunez, A. Rocha, K. Clark, J. M. Sedivy, and N. Neretti, Senescence-associated chromatin rewiring promotes inflammation and transposable element activation, bioRxiv (2025).

[63] E. Schad, P. Tompa, and H. Hegyi, The relationship between proteome size, structural disorder and organism complexity, Genome biology 12, R120 (2011).

