## Supplemental Material for "Chromatin architecture dictates epigenetic memory"

### 1. Detailed derivation of the SWE model

In the following, we derive the spreading-writing-erasing (SWE) model from a discrete model in which the epigenetic mark level of each nucleosome is explicitly modeled:

$$\frac{\partial m_i}{\partial t} = \frac{k_0}{\zeta_0} \sum_j \left[1 - m_i\right] m_j \frac{m_i m_j}{1 + (|i-j|l)^n/\xi^n} - \alpha m_i + \gamma_i. \tag{S1}$$

Here, the summation is over nucleosomes, where $i = 1, 2, \ldots, N$, and $N$ is the total number of nucleosomes. $l$ is the genomic distance between consecutive nucleosomes and $\xi$ is the lower cutoff in the power-law decay of the contact probability. In this work, we set $\xi = 1$ kb for simplicity, without affecting our conclusions. $k_0$ is the total spreading rate summed over the entire chromosome, and the normalization factor $\zeta_0$ is defined as

$$\zeta_0 = \sum_i \sum_j \left[1 - m_i\right] m_j \frac{m_i m_j}{1 + (|i-j|l)^n/\xi^n}. \tag{S2}$$

We convert Equation S1 into a continuous form and introduce $y = il$ such that

$$\frac{\partial m(y)}{\partial t} = \frac{k_0}{\zeta_0 l} \int \left[1 - m(y)\right] m(y') \frac{m(y)m(y')}{1 + |y-y'|^n/\xi^n} dy' - \alpha m(y) + \gamma(y), \tag{S3}$$

and

$$\zeta_0 = \frac{1}{l^2} \int \int \left[1 - m(y)\right] m(y') \frac{m(y)m(y')}{1 + |y-y'|^n/\xi^n} dy' dy. \tag{S4}$$

To obtain the equation in the main text, we introduce the dimensionless coordinate, $x = y/\xi$, such that

$$\frac{\partial m(x)}{\partial t} = \frac{k_0 \xi}{\zeta_0 l} \int \left[1 - m(x)\right] m(x') \frac{m(x)m(x')}{1 + |x-x'|^n} dx' - \alpha m(x) + \gamma(x), \tag{S5}$$

$$\zeta_0 = \frac{\xi^2}{l^2} \int \int \left[1 - m(x)\right] m(x') \frac{m(x)m(x')}{1 + |x-x'|^n} dx' dx = \frac{\xi^2}{l^2} \zeta. \tag{S6}$$

Comparing Equation S5 with Equation 1 in the main text, we find that the coarse-grained spreading rate introduced in the main text, $k$, is related to $k_0$ via $k = k_0 l/\xi$.

### 2. Model with unlimited reader-writer enzymes

In the model with unlimited reader-writer enzymes, we remove the normalization factor of Equation 1 in the main text, and the total mark level is no longer constrained.

$$\frac{\partial m(x,t)}{\partial t} = k \int_0^L \left[1 - m(x)\right] m(x') \frac{m(x)m(x')}{1 + |x-x'|^n}\, dx' - \alpha m(x) + \gamma(x). \tag{S7}$$

Therefore, the compartment dynamics in the unlimited-enzyme model are sensitive to the spreading rate $k$ (Figure S2A-C). In comparison, the SWE model exhibits stable epigenetic memory (Figure S2D). We also apply global disturbance and periodic DNA replication explicitly to both the unlimited-enzyme model (Figure S3A, B) and the SWE model (Figure S3C, D). While the unlimited-enzyme model loses its epigenetic memory quickly, the SWE model recovers perfectly.

#### 3. Derivation of the threshold writing and erasing strengths

We introduce active writing to a nucleation region with size $b$ inside which $\gamma(x) = \gamma_0$ and outside which $\gamma(x) = 0$. Because the nucleation region ($\sim 10^3$ kb) is much smaller than the total chromosome length ($\sim 10^5$ kb), we approximate the rest of the chromosome as a reservoir in a steady state. Consequently, the normalization factor $\zeta$, which is dominated by the rest of the chromosome, remains constant during the initial stage of nucleation. This approach is analogous to methods used in statistical physics, where one can treat the reservoir surrounding a small system as in thermal equilibrium.

Numerical simulations show that the mark level and the spreading rate (see Equation 1 in the main text) are approximately uniform inside compartments (Figure S1). This simplification lets us consider a point near the middle of the compartment to study its mark-level dynamics. By balancing the rates of spreading, active writing, and passive erasing, we find that

$$C(1-m)m^3 = m - \gamma, \tag{S8}$$

with the constant $C = k/\zeta \int_0^{b/2} 2/(1+x^n)dx$. Using the same idea, we derive the equation for the equilibrium mark level upon active erasing, which is Equation 3 in the main text. We confirm Equation 2 and Equation 3 in the main text by simulating the SWE model directly (Figure S4 and Figure S5).

We next study whether the induced compartment can grow beyond the nucleation region. Mathematically, this depends on whether the spreading rate in the nearby unmodified region exceeds the passive erasing strength. From Equation 1 in the main text, we find that for the compartment to grow, the induced mark level must satisfy

$$Dm^2 > 1, \tag{S9}$$

where the constant $D = k/\zeta \int_0^{b} 1/(1+x^n)dx$. Combining Equations S8 and S9, we derive the minimum mark level $m_c$ for the compartment to grow as a function of $\gamma$. The induced mark level always exceeds $m_c$, as long as the size of the nucleation region is not too small (dashed line in Figure 2A of the main text).

#### 4. The value of $k/\zeta$ in Figure 2 of the main text

In Figure 2 of the main text, we focus on a single compartment and treat the rest of the chromosome as a reservoir with fixed $k/\zeta$. We assume the reservoir is composed of $N$ compartments with a typical estimated size $b_0 = 10^{3.5}$ and mark level $m$. Meanwhile, the total mark level in the reservoir is $k$ (considering $\alpha_0 = 1$). Therefore, $k = Nmb_0$ and

$$\frac{k}{\zeta} = \frac{Nmb_0}{N(1-m)m^3}\left[\int_0^{b_0}\int_0^{b_0} \frac{1}{1+|x-x'|^n}dx'dx\right]^{-1} \approx \frac{(1-n)(2-n)}{(1-m)m^2[b_0^{1-n}-(2-n)]}. \tag{S10}$$

We take $m = 0.7$ and $b_0 = 10^{3.5}$ in Equation S10 to calculate $k/\zeta$ as a function of $n$ in Figure 2.

#### 5. The maintenance time as a function of compartment size

In this section, we derive the maintenance time of epigenetic memory and consider a simple case in which the chromosome has two compartments of widths $b+\delta$ and $b-\delta$, respectively. We define the total mark level of a single compartment as $M_i = \int_i m(x)dx$. Here, $\int_i$ represents integration over the region of compartment $i$. The temporal evolution of $M_i$ follows the equation:

$$\frac{\partial}{\partial t}M_i = \frac{k}{\zeta}\zeta_i - M_i. \tag{S11}$$

Here, $\zeta_i$ is defined similarly to $\zeta$, but only integrated within the region of compartment $i$. We approximate the mark level inside all compartments as a constant value $m_0$, consistent with the numerical results presented in Figure 1C of the main text, and therefore $b_i \propto M_i$. We convert Equation S11 into the dynamics of width $b_i$ as

$$\frac{\partial}{\partial t}b_i = \frac{k}{m_0}\frac{\kappa_i}{\sum_j \kappa_j} - b_i, \tag{S12}$$

where

$$\kappa_i = \int_0^{b_i} \int_0^{b_i} \frac{1}{1+|x-x'|^n} dx' dx. \tag{S13}$$

By subtracting the differential equations for $b_1 = b + \delta$ and $b_2 = b - \delta$, we find that the dynamics of the variable $\delta$ satisfies

$$\frac{d\delta}{dt} = \frac{k}{2m_0} \frac{\kappa(b_1) - \kappa(b_2)}{\kappa(b_1) + \kappa(b_2)} - \delta. \tag{S14}$$

If the initial difference is small, $\delta \ll b$, the expression can be expanded around $b = k/(2m_0)$ as

$$\frac{d\delta}{dt} = \left[ b \frac{\kappa'(b)}{\kappa(b)} - 1 \right] \delta. \tag{S15}$$

Thus, we estimate the maintenance time of epigenetic memory as

$$t_m = \left[ \frac{b\kappa'(b)}{\kappa(b)} - 1 \right]^{-1}. \tag{S16}$$

To obtain an expression of maintenance time, we first calculate $\kappa(b)$ as

$$\kappa = \int_0^{b} \int_0^{b} \frac{1}{1+|x-x'|^n} dx' dx = 2 \int_0^b \frac{b-z}{1+z^n} dz. \tag{S17}$$

We can further approximate $\kappa$ as

$$\kappa \approx 2 \int_1^b (b-z) z^{-n} dz \propto \frac{b^{2-n} - b}{1-n} - \frac{b^{2-n} - 1}{2-n}. \tag{S18}$$

In writing the above approximation, we assume $b \gg 1$. Using Equation S16, we find the relationship between $t_m$ and the compartment size $b$ as

$$t_m(b,n) \propto \frac{(2-n)b - b^{2-n}}{(n-1)(b^{2-n} - 1)} \sim \begin{cases} 1 & n < 1 \\ b^{n-1} & 1 < n < 2 \\ b & n > 2. \end{cases} \tag{S19}$$

For $n = 1$, $t_m \sim \ln(b)$ and for $n = 2$, $t_m \sim b/\ln(b)$, both in the large $b$ limit.

### 6. Protocols to enhance reprogramming efficiency

We apply our theory to study iPSC reprogramming efficiency. For simplicity, we simulate a single compartment and treat the rest of the chromosome as a time-independent reservoir. We apply periodic dilution to model DNA replication and monitor the compartment through multiple cell cycles. This process is equivalent to tracking one daughter cell after cell division.

First, we propose that adding noise to the segregation of parental histones during DNA replication may increase reprogramming efficiency (Figure S9A). A sufficiently large fluctuation in parental-histone segregation is more likely to drive the system across the threshold for compartment erasure. To test this hypothesis, we multiply the mark level by a factor of $1/2[1 + \sigma N(0, 1)]$ at each DNA replication event, where $N(0, 1)$ is a standard normal distribution and $\sigma$ is the noise strength. We compute the reprogramming efficiency as the probability that the compartment is successfully erased. In the control case where the mark level is reduced exactly by half during DNA replication, the mark level returns to its target state before each subsequent dilution (the $\sigma = 0$ curve in Figure S9A). Intriguingly, the segregation noise significantly enhances the reprogramming efficiency (Figure S9A). Our results suggest that defects in parental histone recycling, e.g., via MCM2 knockdown, may improve reprogramming efficiency in iPSCs.

Second, we propose that accelerating cell proliferation also increases reprogramming efficiency, since a shorter doubling time is associated with a higher erasing strength in the continuous SWE model. To verify our prediction, at every DNA replication, the time interval to the next DNA replication $T$ follows a random Gaussian distribution value $T = T_0 + \sigma_T N(0, 1)$, while keeping parental histone segregation noiseless. As predicted, a shorter average doubling time than that of wild-type cells results in higher reprogramming efficiency (Figure S9B). Correspondingly, experimental studies have reported that fast-dividing cells lose original epigenetic identity more rapidly than slow-dividing ones because of replicative dilution. Our results also explain why a rapid increase in cell proliferation (e.g., via the c-Myc factor) is one of the most critical hallmarks of successful reprogramming.

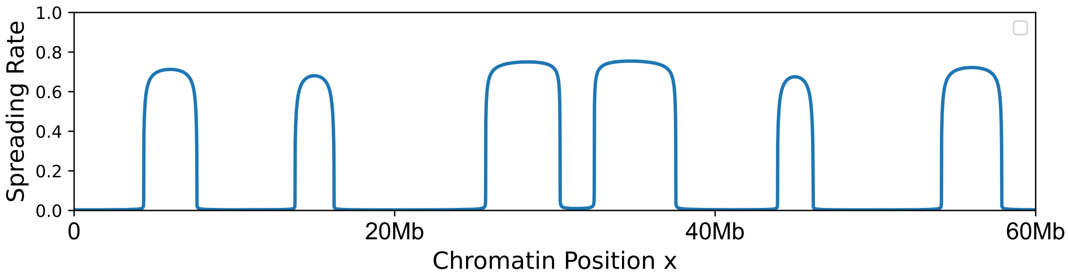


FIG. S1. We calculate the spreading rate, which is $(k/\zeta)\int \frac{[1-m(x)]m(x)m(x')^2}{1+|x-x'|^n}dx'$ from the same simulation as Figure 1B in the main text at generation 0. We note that the spreading rate is approximately uniform inside compartments.

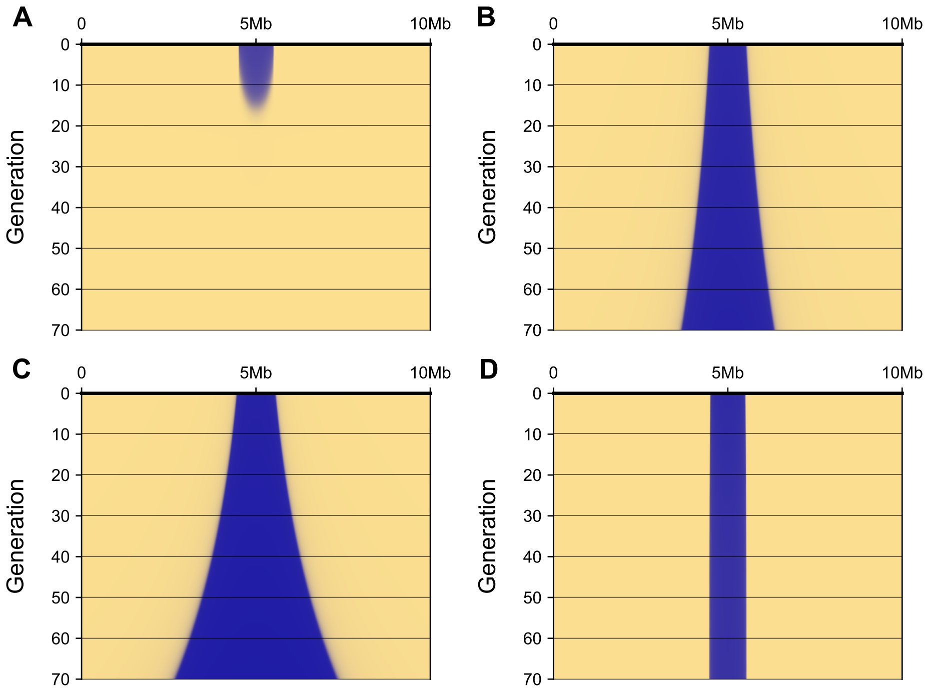


FIG. S2. Simulations of the model with unlimited reader-writer enzymes, governed by Equation S7. Here, we show (A) $k = 7 \times 10^{-3}$, (B) $k = 8 \times 10^{-3}$ and (C) $k = 9 \times 10^{-3}$. The compartment either expands indefinitely or vanishes rapidly, indicating the absence of stable epigenetic memory. For comparison, we also simulate the SWE model with identical initial states in (D), which exhibits stable epigenetic memory.

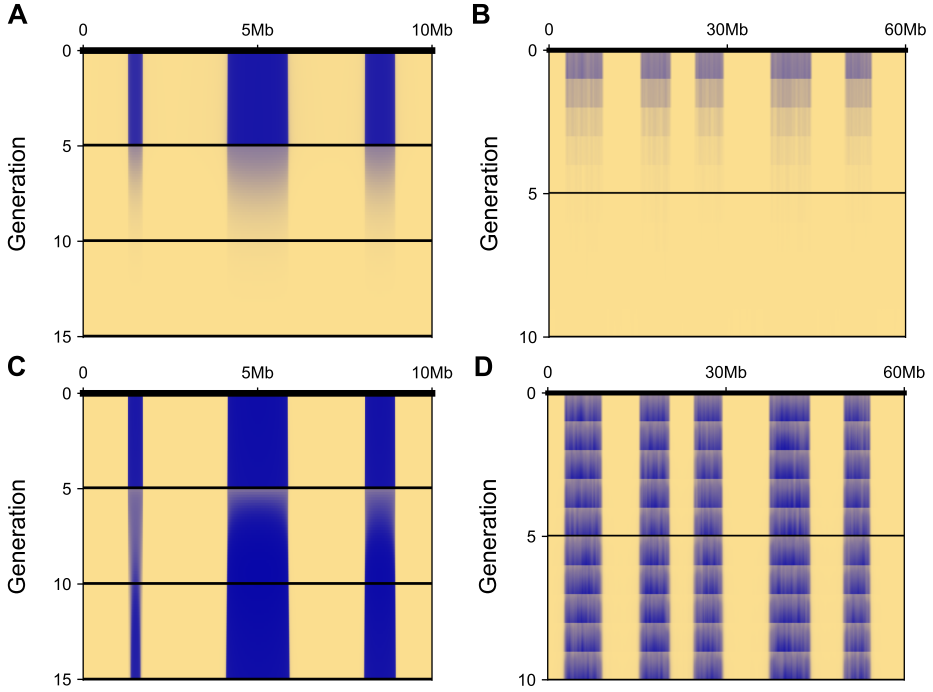


FIG. S3. In the model with unlimited reader-writer enzymes, epigenetic marks are readily erased by strong disturbance or explicit DNA replication in (A) and (B). For comparison, we also simulate the SWE model with identical initial states in (C) and (D). In (A) and (C), we multiply the mark level by $\beta = 0.5$ at $T = 5$. In (B) and (D), we include $\alpha_b = 0.3$ as the basal dilution rate.

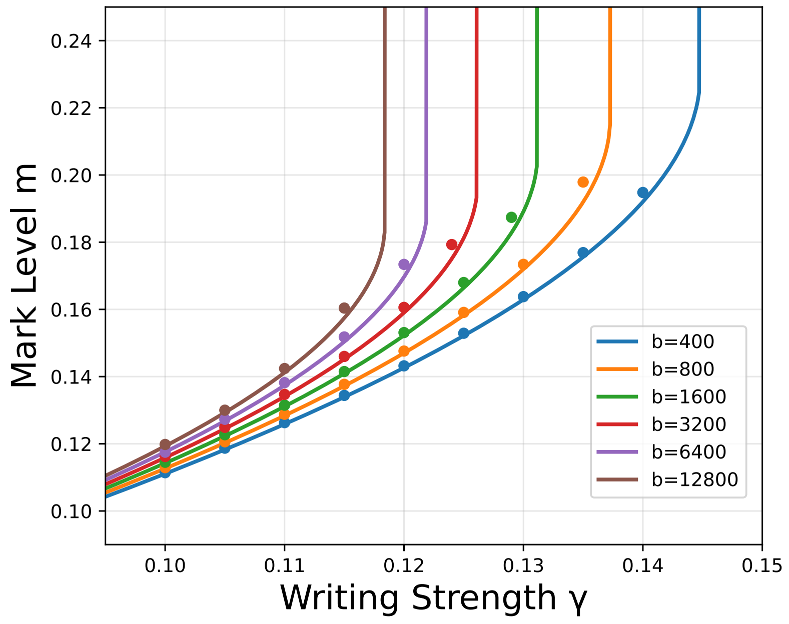


FIG. S4. We validate the approximation made in Equation 2 in the main text by simulating a subsystem embedded in a reservoir with $k/\zeta = 1.1$. The simulation results (dots) agree well with the theoretical prediction (lines).

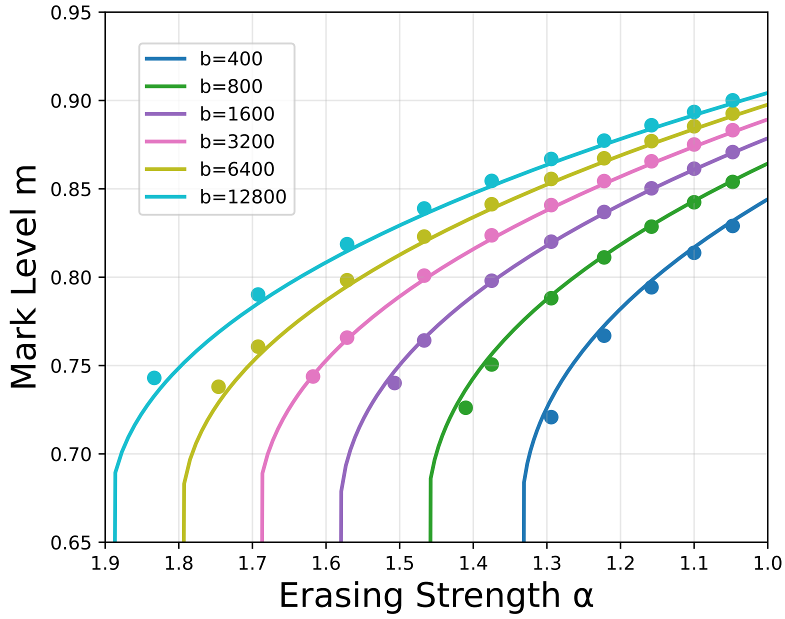


FIG. S5. We validate the approximation made in Equation 3 in the main text by simulating a subsystem embedded in a reservoir with $k/\zeta = 1.1$. The simulation results (dots) agree well with the theoretical prediction (lines).

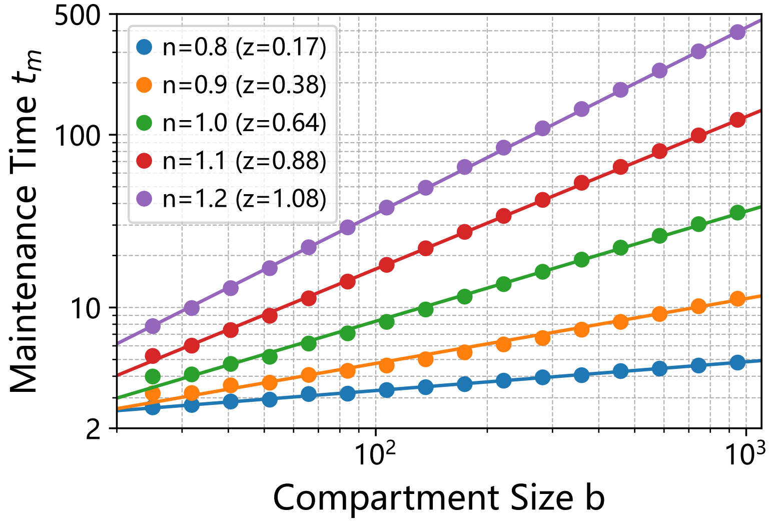


FIG. S6. An approximate power-law scaling $t_m \propto b^z$ exists between the epigenetic-memory maintenance time $t_m$ and the compartment size $b$. The solid lines are power-law fits to the data.

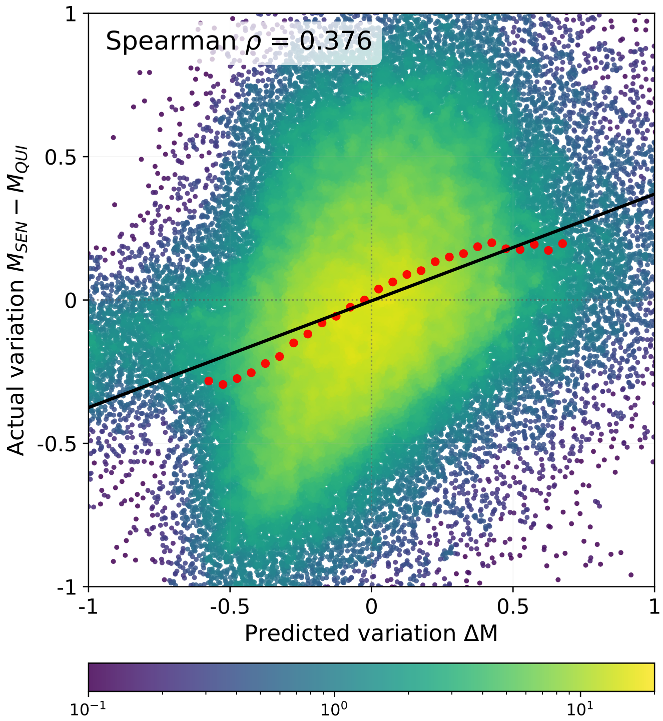


FIG. S7. Same analysis as in Figure 3D of the main text, but for quiescent and senescent cells.

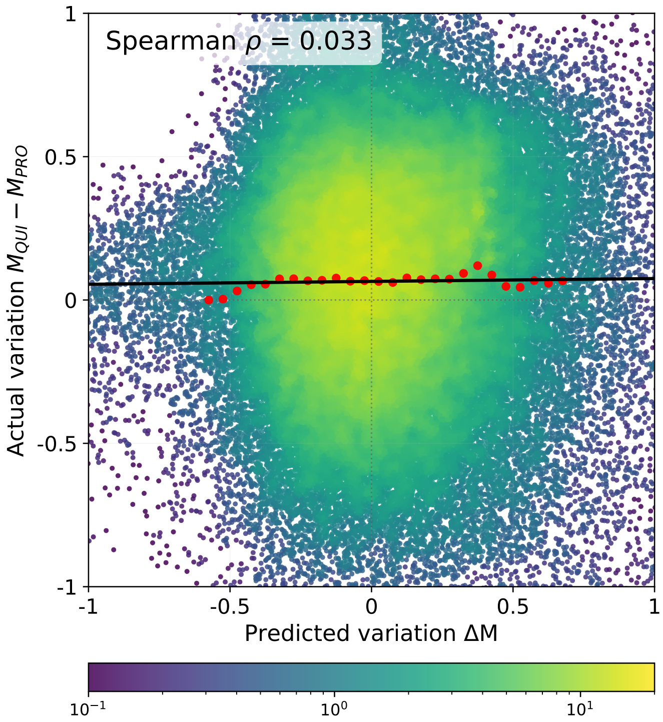


FIG. S8. Same analysis as in Figure 3D of the main text, but for proliferative and quiescent cells.

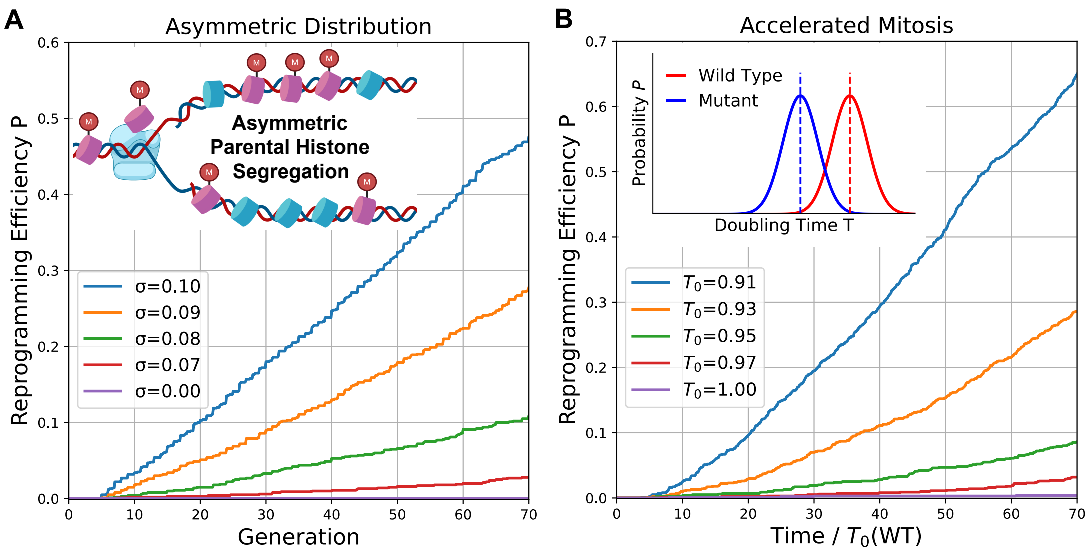


FIG. S9. Protocols to enhance iPSC reprogramming efficiency. (A) Increasing the noise strength $\sigma$ in parental histone segregation increases reprogramming efficiency. (B) Shortening the mean doubling time $T_0$ increases reprogramming efficiency, with $\sigma_T = 0.1$. The x-axis is the time normalized by the mean doubling time of the wild-type cells as $T_0(\mathrm{WT}) = 1$. Inset: the normal distributions of doubling time for wild-type (red) and mutant (blue) cells. Here, we run $10^3$ independent simulations for each case and compute the reprogramming efficiency as the fraction of simulations in which the compartment is erased. In both panels, we take $k/\zeta = 0.8$, $b = 500$, and $\alpha_b = 0.3$ as the basal dilution rate.
